# Spatial structure governs the mode of tumour evolution

**DOI:** 10.1101/586735

**Authors:** Robert Noble, Dominik Burri, Jakob Nikolas Kather, Niko Beerenwinkel

## Abstract

Characterizing the mode – the way, manner, or pattern – of evolution in tumours is important for clinical forecasting and optimizing cancer treatment. DNA sequencing studies have inferred various modes, including branching, punctuated and neutral evolution, but it is unclear why a particular pattern predominates in any given tumour.^1, 2^ Here we propose that differences in tumour architecture alone can explain the variety of observed patterns. We examine this hypothesis using spatially explicit population genetic models and demonstrate that, within biologically relevant parameter ranges, human tumours are expected to exhibit four distinct onco-evolutionary modes (oncoevotypes): rapid clonal expansion (predicted in leukaemia); progressive diversification (in colorectal adenomas and early-stage colorectal carcinomas); branching evolution (in invasive glandular tumours); and effectively almost neutral evolution (in certain non-glandular and poorly differentiated solid tumours). We thus provide a simple, mechanistic explanation for a wide range of empirical observations. Oncoevotypes are governed by the mode of cell dispersal and the range of cell-cell interaction, which we show are essential factors in accurately characterizing, forecasting and controlling tumour evolution.

---

A tumour is a product of somatic evolution in which mutation, selection, genetic drift, and cell dispersal generate a patchwork of cell subpopulations (clones) with varying degrees of aggressiveness and treatment sensitivity.^3^ A primary goal of modern cancer research is to characterize this evolutionary process, to enable precise, patient-specific prognostic forecasts and to optimize targeted therapy regimens. However, studies revealing the evolutionary features of particular cancers raise as many questions as they answer. Why do different tumour types exhibit different modes of evolution^1, 2, 4–8^? What conditions sustain the frequently observed pattern of branching evolution, in which clones diverge and evolve in parallel^1, 9^? And why do some pan-cancer analyses indicate that many tumours evolve neutrally,^10^ whereas others support extensive selection^11^?

Factors proposed as contributing to tumour evolution include microenvironmental heterogeneity, niche construction, and positive ecological interactions between clones.^3, 12, 13^ However, because such factors have not been well characterized across human cancer types, it remains unclear how they might relate to evolutionary modes. On the other hand, it is well established that tumours exhibit a wide range of architectures (Figure 1), the evolutionary effects of which have not been systematically examined. Because gene flow is among the principle forces that shape the genetic makeup of a population, we hypothesized that different tumour structures might result in different tumour evolutionary modes. To test this hypothesis, we developed a way to formulate multiple classes of mathematical models, each tailored to a different class of tumour, within a single general framework, and we implemented this framework as a stochastic computer program.

**Figure 1:**
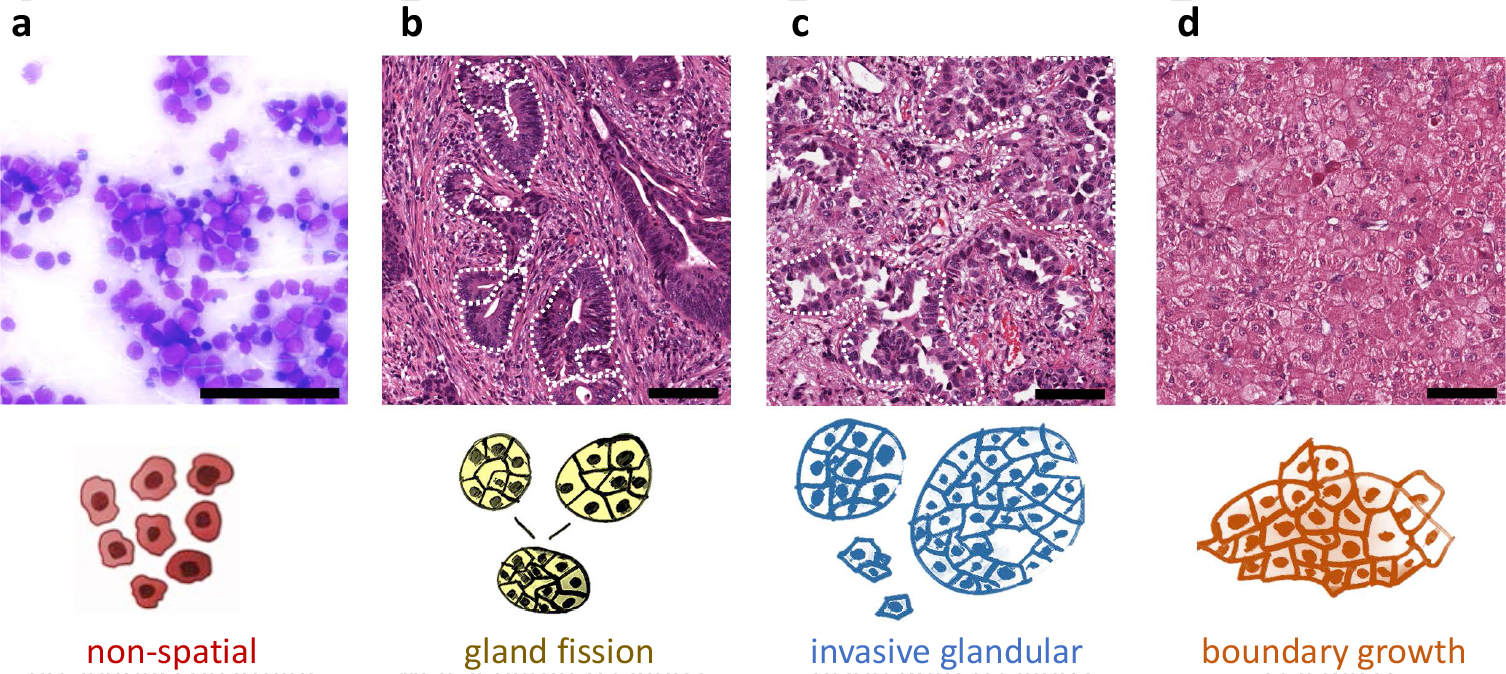
Representative regions from histopathological slides from human tumours exemplifying four different kinds of tissue structure and modes of cell dispersal. **a**, Acute myeloid leukaemia, M2 subtype, bone marrow smear. **b**, Early stage colon cancer (patient TCGA-A6-2684, slide 01Z-00-DX1). **c**, Non-small cell lung cancer (patient TCGA-44-6147, slide 01Z-00-DX1). **d**, Hepatocellular carcinoma (patient TCGA-CC-5258, slide 01Z-00-DX1). Image a is courtesy of Cleo-Aron Weis; images b, c and d were retrieved from The Cancer Genome Atlas at https://portal.gdc.cancer.gov, with brightness and contrast adjusted linearly for better visibility. Scale bars are 100 *μ*m. Dotted curves outline exemplary tumour glands (b and c).

Our approach is built on basic tenets of cancer evolutionary theory.^3^ Simulated tumours arise from a single cell that has acquired a fitness-enhancing mutation. Each time a tumour cell divides, its daughter cells can acquire passenger mutations, which have no fitness effect, and more rarely driver mutations, which confer a fitness advantage. In solid tumours, we assume that cells compete with one another for space and other resources. Whereas previous studies have assumed that tumours grow into empty space, our model also allows us to simulate the invasion of normal tissue that is a defining feature of malignancy. To recapitulate different tumour architectures, we vary parameters of gland size and the mode of cell dispersal, while keeping constant the driver mutation rate and the distribution of driver mutation-induced fitness effects. All tumours take a similar amount of time to grow from one cell to one million cells, corresponding to several years in real time.

Our first case is a non-spatial model that has been proposed as appropriate to leukaemia,^14^ a tumour type in which mutated stem cells in semi-solid bone marrow produce cancer cells that mix and proliferate in the bloodstream. In the absence of spatial constraints, rapid clonal expansions can result from driver mutations that increase the cell division rate by as little as a few percent. For plausible parameter values, the vast majority of cells at the end of the simulation share the same set of driver mutations (Figure 2a-d). These dynamics indeed resemble those of blood cancers such as chronic myeloid leukaemia, which are the least spatially structured of human tumours, and in which cell proliferation is driven by a single change to the genome.^15^

**Figure 2:**
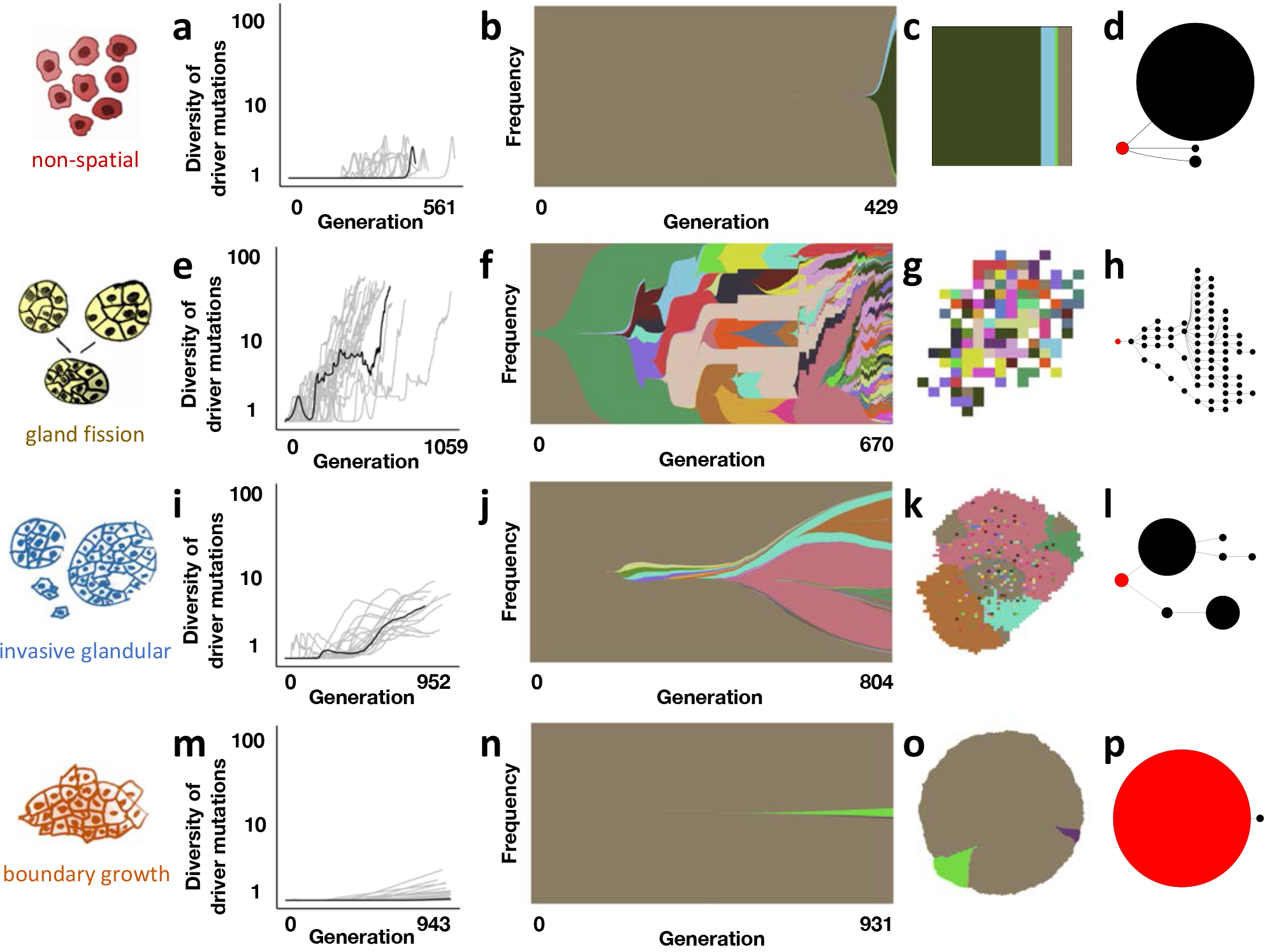
Four oncoevotypes predicted by our model. **a**, Dynamics of driver mutation diversity in 20 stochastic simulations of a non-spatial branching process (initial death rate 0.98, relative to initial division rate). Diversity corresponds to the number of clones that have distinct combinations of driver mutations. A generation is defined as the expected cell cycle time of the initial tumour cell. Black curves correspond to the individual simulations illustrated in subsequent panels (having metrics closest to the medians of sets of 100 replicates). **b**, Muller plots of clonal dynamics over time, for a non-spatial branching process. Colours represent clones with distinct combinations of driver mutations (the original clone is grey-brown; subsequent clones are coloured using a recycled palette of 26 colours). Descendant clones are shown emerging from inside their parents. **c**, Final clone proportions. **d**, Driver phylogenetic trees. Node size corresponds to clone population size at the final time point and the founding clone is coloured red. Only clones whose descendants represent at least 1% of the final population are shown. **e-h**, Results of a model of tumour growth via gland fission (8,192 cells per gland). In the spatial plot (g), each pixel corresponds to a patch of cells, corresponding to a tumour gland, coloured according to the most abundant clone within the patch. **i-l**, Results of a model in which tumour cells invade normal tissue at the tumour boundary (512 cells per gland). **m-p**, Results of a boundary-growth model of a non-glandular tumour. In all cases, the driver mutation rate is 10^−5^ per cell division, and the mean of the exponential distribution of driver fitness effects is 10%. Other parameter values are listed in Supplementary Tables 1 and 2. Muller plots were drawn using the ggmuller R package.^16^

In our second model, consistent with the biology of colorectal adenomas and early-stage colorectal carcinomas,^17^ and in common with previous studies,^5, 18^ we simulate a tumour that consists of large glands and that grows via gland fission (bifurcation). Although the driver mutation rate and the fitness effect are exactly the same as in the previous case, the addition of spatial structure dramatically alters the evolutionary mode. The organization of cells into glands limits the extent to which driver mutations can spread through the population. As the tumour grows larger, selective sweeps become progressively localized, leading to a fan-like driver phylogenetic tree and ever greater spatial diversity, with different combinations of driver mutations predominating even in neighbouring glands (Figure 2e-h). This pattern is maintained even if cells are able to acquire drivers that directly increase the gland fission rate, because such mutations rarely spread within glands (Supplementary figure 1a).

Our third case corresponds to a glandular (acinar) tumour that grows by tumour budding and invasion of normal tissue. At later stages, solid tumours of many types – including most well-differentiated adenocarcinomas and squamous cell carcinomas of multiple primary sites – assume such a structure. To parameterize an appropriate model, we used semi-automated analysis of histopathological slides (Supplementary figure 2) and found that each gland of an invasive, acinar tumour can contain between a few hundred and a few thousand cells (Supplementary Table 3). In contrast to the fission case, simulated invasive glandular tumours typically exhibit stepwise increases in driver diversity and a phylogeny with multiple long branches (Figure 2i-l). Because competition between tumour cells and normal cells amplifies selection, even small increases in cell fitness can spark rapid clonal expansions. Nevertheless, even if cells are able to invade neighbouring glands within the tumour bulk (Supplementary figure 1b), clonal interference inhibits selective sweeps. The result is a zonal tumour, with large regions sharing the same combination of driver mutations. Branching evolution has indeed been associated with invasive glandular morphology in numerous cancer types.^1, 2^

Our fourth and final model represents a tumour with no glandular structure and with growth confined to its boundary. Tumour types such as hepatocellular carcinoma, undifferentiated carcinomas of multiple primary sites, and some benign tumours frequently comprise a dense mass of cells with a well-delineated boundary around each tumour nodule. Accordingly, the boundary growth model has been proposed as particularly appropriate for simulating the evolution of hepatocellular carcinoma nodules.^7, 19^ The spatial structure of this model favours genetic drift, rather than selection. For our fixed parameter values, tumour evolution in the boundary-growth case is effectively almost neutral (Figure 2m-p), and mutations can spread only by “surfing” on a wave of population expansion.^20–22^ Selection is only slightly more prominent when cells can compete with neighbours within the tumour mass (Supplementary figure 1c). Such suppression of selection in the boundary-growth model is consistent with evidence of effectively neutral evolution in hepatocellular carcinoma,^7^ as well as the existence of large, non-glandular benign tumours that only rarely progress to malignancy.

For any given tumour architecture, the evolutionary mode is predicted to shift towards the effectively neutral case when the mutation rate is lower, the driver fitness effect is smaller, or initial tumour growth is very rapid, relative to the cell division rate. Our framework also implies that a change in tumour architecture during cancer progression will lead to a change in oncoevotype. For example, in contrast to the “big bang” model of colorectal cancer,^4, 5^ we predict ongoing selection in colorectal adenomas (whose growth is driven by gland fission), enabling multiple driver mutations to reach high frequencies. After an invasive subclone of the adenoma gives rise to a carcinoma, we predict a transition to either branching evolution or – because the invasion begins with a highly transformed, rapidly expanding subclone – effectively neutral evolution. This explanation is broadly consistent with recent multi-region sequencing studies,^6, 23^ while also agreeing with results of comparative genomic analysis, which indicate that colorectal cancers evolve subject to strong positive selection and have more driver mutations per cell than most other cancer types.^11^

Overall, our models demonstrate that variation in the range of cell-cell interaction and the mode of cell dispersal is sufficient to generate distinct tumour evolutionary modes or oncoevotypes (Table 1). These oncoevotypes can moreover be clearly distinguished using two simple, intuitive measures. The first metric is the diversity of driver mutations, which roughly corresponds to the breadth of the driver phylogenetic tree (as in the final column of Figure 2). The second metric is the mean number of driver mutations per cell, which represents the depth of the driver phylogenetic tree. In terms of these evolutionary metrics, the four oncoevotypes discussed above form four distinct clusters (Figure 3a). Neutral counterparts of these four models – which have identical parameter values, except that drivers have no phenotypic effect – cluster together, near the boundary-growth model.

**Table 1:** Properties of four archetypal oncoevotypes during initial tumour growth.

| Oncoevotype | Driver diversity | Drivers per cell | Role of selection | Associated tumour characteristics |
| --- | --- | --- | --- | --- |
| Selective sweeps | Low | Low | Strong | Little spatial structure |
| Progressive diversification | High | High | Strong only at the local level | Gland fission |
| Deep branching | Intermediate | Intermediate | Strong, but constrained by clonal interference and/or microenvironmental factors | Glandular; tumour budding; infiltration |
| Effectively almost neutral | Low | Low | Weak | Expansive growth; rapid growth |

**Figure 3:**
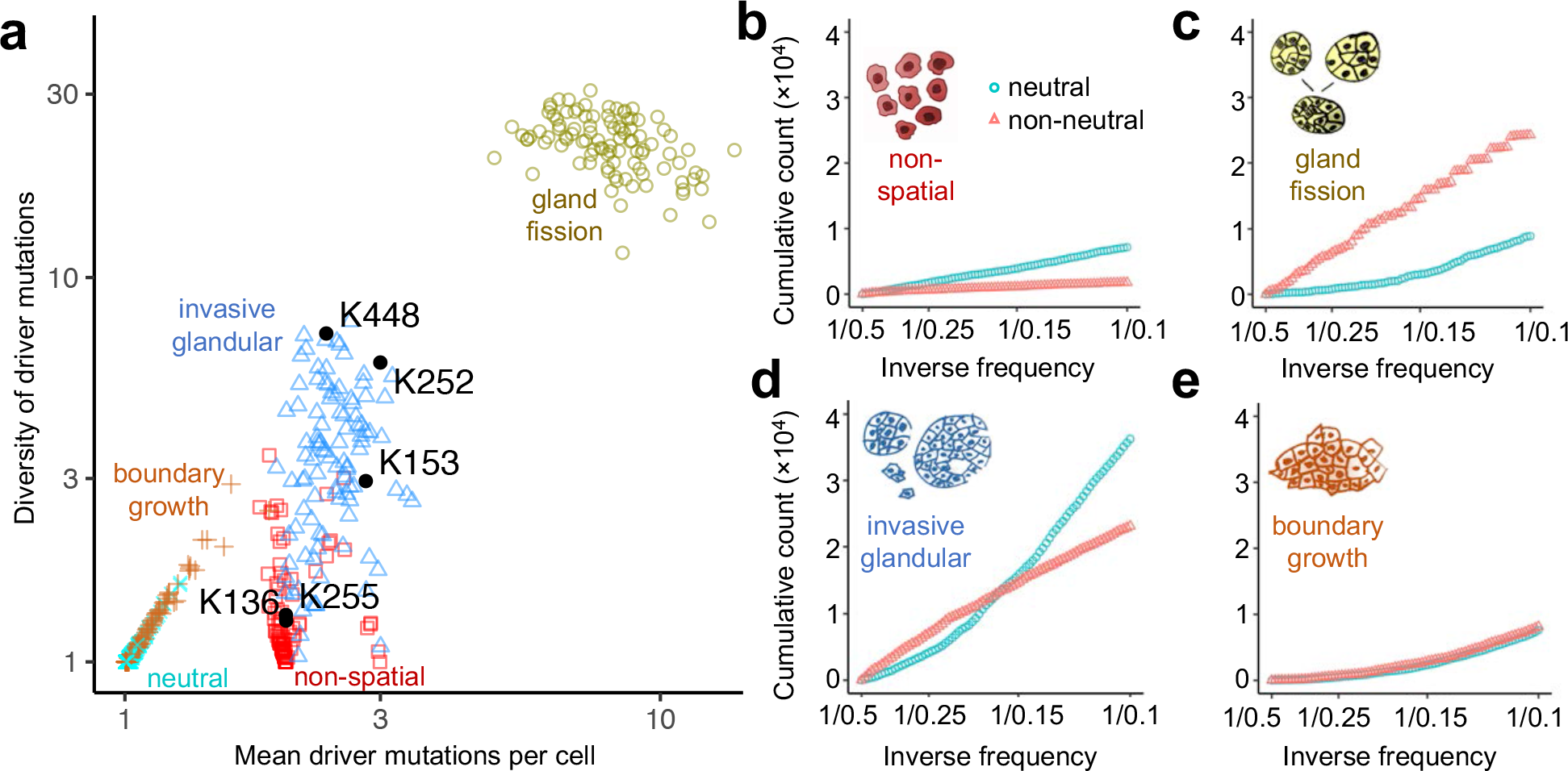
Oncoevotypes distinguished in terms of summary metrics and mutation frequencies. **a**, Evolutionary metrics of four example models with different spatial structures and different modes of cell dispersal but identical driver mutation rates and identical driver mutation effects (100 stochastic simulations per model). Neutral counterparts of the four models are represented together as an additional group. Black points correspond to kidney tumour (ccRCC) data, labelled with patient codes.^9^ **b-e**, Mutant allele frequency distributions predicted by our model for simulations with only neutral mutations (blue points) or both neutral and driver mutations (red points). Cumulative mutant allele count is plotted against inverse mutant allele frequency (1/*f*), restricted to mutations with frequencies between 0.1 and 0.5. Each distribution represents combined data from 100 simulations. Parameter values for the four models are the same as in Figure 2.

Few studies have examined tumour evolutionary trajectories in sufficient detail to enable quantitative comparison with our model results. The most remarkable example is a recent multi-centre study of clear cell renal cell carcinoma (ccRCC) that involved multi-region, deep sequencing of 101 tumours, targeting a panel of more than 100 putative driver genes.^9^ This data set is ideally suited for generating driver phylogenetic trees that are readily comparable to those predicted by our modelling. We focus on five cases of ccRCC for which driver frequencies were reported in the original publication, and which are representative of the larger cohort in exhibiting branching evolution, with between three and nineteen driver mutations per tumour. Since ccRCC is typically an invasive glandular tumour, our framework predicts that ccRCC should exhibit branching evolution. Indeed, we find that the evolutionary metrics and driver phylogenetic tree structures of the five ccRCC tumours are highly consistent with the predicted oncoevotype (Figure 3a; Supplementary figure 3). This result is robust to varying gland size within plausible ranges (Supplementary figures 1d and 4a).

Because researchers and clinicians seldom have access to multi-regional sequencing data, nor the longitudinal data needed to track how tumour clone sizes change over time, tumour phylogenies and evolutionary parameters are more usually inferred from mutation frequencies measured from a single biopsy sample at a single time point. Moreover, current cancer sequencing technologies are neither sensitive enough to detect the majority of low frequency mutations, nor precise enough to distinguish between high frequency and clonal (100% frequency) mutations. Accordingly, the most relevant part of the mutation frequency distribution for practical purposes is in the intermediate frequency range. One way to examine differences between distributions within this intermediate range is to plot the cumulative mutation count (the number of mutations present at or above frequency *f*) versus the inverse mutation frequency (1/*f*). In a neutral non-spatial model, this graph is a straight line (Figure 3d, blue points). The most compelling evidence for widespread neutral evolution in human cancers is based on the observation that the transformed mutation frequency distributions of many tumours are also approximately linear.^10^

Our population genetic modelling indicates that tumour architecture has important effects on tumour mutation frequency distributions (Figure 3e-g; Supplementary figure 5). In particular, when the cumulative mutation count is plotted against the inverse mutation frequency, the curve for the neutral model is no longer linear. On the contrary, the average non-neutral model curve can be closer to a straight line than the average neutral model curve. It follows that methods using mutation frequencies to infer selection in solid tumours will yield incorrect conclusions if they fail to account for effects of population structure. Inappropriate choice of null model can therefore explain otherwise contradictory findings regarding the prevalence of neutral evolution in human cancers.^11, 24^

In summary, we have found that differences in the range of cell-cell interaction and the mode of cell dispersal can explain the spectrum of evolutionary modes observed in human tumours. Whereas previous mathematical modelling studies have focussed on fitness effects of driver mutations,^5, 10, 18, 25, 26^ our perspective instead emphasizes the importance of population structure and gene flow in tumour evolution. It follows that tumour architecture determines how well biopsy samples reflect intra-tumour heterogeneity. Oncologists typically base treatment decisions on the presence or absence of particular mutations in cells taken from only a small region of a solid tumour. In general, our models predict that biopsies will be most representative of non-glandular tumours with well-delineated boundaries, such as hepatocellular carcinoma nodules, and least representative of tumours that grow via glandular fission, such as early-stage colorectal carcinomas (Figure 2). Tumour types with structures that promote diversification are predicted to be the least responsive to targeted therapies unless truncal mutations can be reliably identified and targeted.

In clear cell renal cell carcinoma, separate studies have found that tumour architecture^27, 28^ and evolutionary trajectory^9^ are predictors of cancer progression and survival. Evolutionary mode also correlates with both tumour architecture and clinical outcome in childhood cancers.^8^ By mechanistically connecting tumour architecture to oncoevotype, our work provides the blueprint for a new generation of patient-specific models for forecasting tumour progression and for optimizing treatment regimens that exploit evolutionary dynamics.^29, 30^

## Data availability

Raw output files from the computational models are available on request.

## Code availability

Computational modelling code is available in an online repository.^31^

## Acknowledgments

We thank Mykola Lebid, Katharina Jahn, Richard Neher, Andreas Deutsch, Kiril Korolev, Cleo-Aron Weis, Benjamin Werner and Andrea Sottoriva for helpful discussions. R.N. and N.B. were supported by ERC Synergy Grant 609883. J.N.K. was supported by the German Consortium for Translational Cancer Research (DKTK/DKFZ) fellowship program and by RWTH Aachen START grant 2018/691906. R.N. was also supported by the National Cancer Institute of the National Institutes of Health under Award Number U54CA217376. The content is solely the responsibility of the authors and does not necessarily represent the official views of the National Institutes of Health.

## Author contributions

R.N. conceived the research question and designed and created the modelling framework. R.N. and D.B. performed analysis of data from the model. J.N.K. obtained and analysed histopathological data. R.N. wrote the manuscript with critical comments and input from N.B., D.B. and J.N.K. All authors have read and edited the final manuscript.

## Methods

### Previous mathematical models of tumour population genetics

Many previous studies of tumour population genetics have used non-spatial branching processes,^14, 25^ in which cancer clones grow exponentially without interacting. Among spatial models, a popular option is the Eden growth model (or boundary-growth model), in which cells are located on a regular grid with a maximum of one cell per site, and a cell can divide only if an unoccupied neighbouring site is available to receive the new daughter cell.^19, 32^ Other methods with one cell per site include the voter model^19, 33, 34^ (in which cells can invade neighbouring occupied sites) and the spatial branching process (in which cells budge each other to make space to divide). Further mathematical models have been designed to recapitulate glandular tumour structure by allowing each grid site or “deme” to contain multiple cells and by simulating tumour growth via deme fission throughout the tumour^5^ or only at the tumour boundary.^18^ A class of model in which cancer cells are organized into demes and disperse into empty space has also been proposed.^22, 35^ Supplementary Table 4 summarizes selected studies representing the state of the art of stochastic modelling of tumour population genetics.

Our main methodological innovations are to implement all of these distinct model structures, as well as models of invasive tumours, within a common framework, and to combine them with methods for tracking driver and passenger mutations at single-cell resolution. The result is a highly flexible framework for modelling tumour population genetics that can be used to examine consequences of variation not only in mutation rates and selection coefficients, but also in factors that control gene flow.^31^

### Computational model structure

Simulated tumours in our models are made up of patches of interacting cells located on a regular grid of sites. In keeping with the population genetics literature, we refer to these patches as demes. All demes within a model have the same carrying capacity, which can be set to any positive integer. Each cell belongs to both a deme and a genotype. If two cells belong to the same deme and the same genotype then they are identical in every respect, and hence the model state is recorded in terms of such subpopulations rather than in terms of individual cells. For the sake of simplicity, computational efficiency, and mathematical tractability, we assume that cells within a deme form a well-mixed population. The well-mixed assumption is consistent with previous mathematical models of tumour evolution^5, 18, 22, 35^ and with experimental evidence in the case of stem cells within colonic crypts.^36^

### Initial conditions

A simulation begins with a single tumour cell located in a deme at the centre of the grid. If the model is parametrized to include normal cells then these are initially distributed throughout the grid such that each deme’s population size is equal to its carrying capacity. Otherwise, if normal cells are absent, then the demes surrounding the tumour are initially unoccupied.

### Stopping condition

The simulation stops when the number of tumour cells reaches a threshold value. Because we are interested only in tumours that reach a large size, if the tumour cell population succumbs to stochastic extinction then results are discarded and the simulation is restarted (with a different seed for the pseudo-random number generator).

### Within-deme dynamics

Tumour cells undergo stochastic division, death, dispersal, and mutation events, whereas normal cells undergo only division and death. The within-deme death rate is density-dependent. When the deme population size is less than or equal to the carrying capacity, the death rate takes a fixed value *d*_0_ that is less than the initial division rate. When the deme population size exceeds carrying capacity, the death rate takes a different fixed value *d*_1_ that is much greater than the largest attainable division rate. Hence all genotypes grow approximately exponentially until the carrying capacity is attained, after which point the within-deme dynamics resemble a birth-death Moran process – a standard, well characterized model of population genetics.

In all spatially structured simulations we set *d*_0_ = 0 to prevent demes becoming empty. For the well-mixed model, we set *d*_0_ > 0 and dispersal rate equal to zero, so that all cells always belong to a single deme (with carrying capacity greater than the maximum tumour population size).

### Mutation

When a cell divides, each daughter cell inherits its parent’s genotype plus a number of additional mutations, drawn from a Poisson distribution. Each mutation is unique, consistent with the infinite-sites assumption of canonical population genetics models. Whereas some previous studies have examined the effects of only a single driver mutation (Supplementary Table 4), in our model there is no limit on the number of mutations a cell can acquire. Most mutations are passenger mutations with no phenotypic effect. The remainder are drivers, each of which increases the cell division or dispersal rate.

The program records the immediate ancestor of each clone (defined in terms of driver mutations) and the matrix of hamming distances between clones (that is, how many driver mutations are not shared by each pair of clones), which together allow us to reconstruct driver phylogenetic trees. To improve efficiency, the distance matrix excludes clones that failed to grow to more than ten cells and failed to produce any other clone before becoming extinct.

### Driver mutation effects

Whereas previous models have typically assumed that the effects of driver mutations combine multiplicatively, this can potentially result in implausible trait values (especially in the case of division rate if the rate of acquiring drivers scales with the division rate). To remain biologically realistic, our model invokes diminishing returns epistasis. Specifically, the effect of a driver is to multiply the trait value *r* by a factor of 1 + *s*(1 − *r*/*m*), where *s* > 0 is the mutation effect and *m* is an upper bound. Nevertheless, because we set *m* to be much larger than the initial value of *r*, the combined effect of drivers in all models in the current study is approximately multiplicative. For each mutation, the value of the selection coefficient *s* is drawn from an exponential distribution.

### Dispersal

Depending on model parametrization, dispersal occurs via either invasion or deme fission (Supplementary Table 1). In the case of invasion, the dispersal rate corresponds to the probability that a cell newly created by a division event will immediately attempt to invade a neighbouring deme. This particular formulation ensures consistency with a standard population genetics model known as the spatial Moran process. Because the dispersal rate is a probability, all values greater than or equal to 1 are equivalent. The destination deme is chosen uniformly at random from the four nearest neighbours (Von Neumann neighbourhood). Invasion can be restricted to the tumour boundary, in which case the probability that a deme can be invaded is *N*/*K*, where *N* is the number of tumour cells in the deme and *K* is the carrying capacity. If a cell fails in an invasion attempt then it remains in its original deme. If invasion is not restricted to the tumour boundary then invasion attempts are always successful.

In fission models, a deme can undergo fission only if its population size is greater than or equal to carrying capacity. As with invasion, deme fission immediately follows cell division (so that results for the different dispersal types are readily comparable). The probability that a deme will attempt fission is equal to the sum of the dispersal rates of its constituent cells. Deme fission involves moving half of the cells from the original deme into a new deme, which is placed beside the original deme. If the dividing deme contains an odd number of cells then the split is necessarily unequal, in which case each deme has a 50% chance of receiving the larger share. Genotypes are redistributed between the two demes without bias according to a multinomial distribution. Cell division rate has only a minor effect on deme fission rate because a deme created by fission takes only a single cell generation to attain carrying capacity.

If fission is restricted to the tumour boundary then the new deme’s assigned location is chosen uniformly at random from the four nearest neighbours, and if the assigned location already contains tumour cells then the fission attempt fails. If fission is allowed throughout the tumour then an angle is chosen uniformly at random, and demes are budged along a straight line at that angle to make space for the new deme beside the original deme.

Our particular method of cell dispersal is chosen to enable comparison between our results and those of previous studies and to facilitate mathematical analysis. In particular, when the deme carrying capacity is set to 1, our model approximates an Eden growth model (if fission is restricted to the tumour boundary, or if dispersal is restricted to the tumour boundary and normal cells are absent), a voter model (if invasion is allowed throughout the tumour), or a spatial branching process (if fission is allowed throughout).

To fairly compare different spatial structures and modes of cell dispersal, we set dispersal rates in each case such that the time taken for a tumour to grow from one cell to one million cells is approximately the same as in the neutral Eden growth model with maximal dispersal rate. This means that, across models, the cell dispersal rate decreases with increasing deme size. Given that tumour cell cycle times are of the order of a few days, the timespans of several hundred cell generations in our models realistically correspond to several years of tumour growth.

### Two versus three dimensions

We chose to conduct our study in two dimensions for two main reasons. First, the effects of deme carrying capacity on evolutionary dynamics are qualitatively similar in two and three dimensions, yet a two-dimensional model is simpler, easier to analyse, and easier to visualize. Second, we aimed to create a method that is readily reproducible using modest computational resources and yet can represent the long-term evolution of a reasonably large tumour at single-cell resolution.

One million cells in two dimensions corresponds to a cross-section of a three-dimensional tumour with many more than one million cells. Therefore, compared to a three-dimensional model, a two-dimensional model can provide richer insight into how evolutionary dynamics change over time. Developing an approximate, coarsegrained analogue of our model that can efficiently simulate the population dynamics of very large tumours with different spatial structures in three dimensions is an important direction for future research.

### Implementation

The program implements Gillespie’s exact stochastic simulation algorithm^37^ for statistically correct simulation of cell events. The order of event selection is 1. deme, 2. cell type (normal or tumour), 3. genotype, and 4. event type. At each stage, the probability of selecting an item (deme, cell type, genotype or event type) is proportional to the sum of event rates for that item, within the previous item.

### Diversity metric

To measure diversity of driver mutation combinations, we use the inverse Simpson index defined as 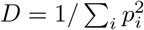, where *p*_*i*_ is the frequency of the *i*th combination of driver mutations. For example, if the population comprises *k* types of equal size then *p*_*i*_ = 1/*k* for every value of *i*, and so *D* = 1/(*k* × 1/*k*^2^) = *k*. Furthermore, in the hypothetical case where each clone is replaced by *α* daughter clones with equal frequencies, we have *D* = 2^*α*(*n*−1)^, where *n* is the average number of drivers per cell.

Our diversity metric fulfils the same purpose as the intratumour heterogeneity (ITH) measure used in the TRACERx Renal study,^9^ and indeed the two metrics are strongly correlated across our models (Spearman’s *ρ* = 0.93). Our metric has three main advantages compared to ITH: first, *D* is a continuous variable; second, *D* is robust to methodological differences that affect ability to detect low frequency mutations; third, *D* is not directly dependent on our second evolutionary metric (the number of driver mutations per cell).

### Histopathological slide analysis

We estimated the number of cells per gland in well-differentiated adenocarcinomas – one of the most abundant types of human cancer across primary sites – using semi-automated analysis of histopathological slides selected at random for three patients with advanced colorectal cancer, a tumour type with pronounced stromal infiltration and spatially separated glands.^38^ We delineated a total of 41 clearly separated glands in whole-slide images from the three patients and counted the cells in each gland with digital pathology software^39^ (Supplementary figure 2).

In cross section, the number of cells per gland ranged from 14 to 1,563, with 68% of cases between 50 and 500 cells (Supplementary Table 3). It is therefore reasonable to assume that each gland of an invasive, acinar tumour can contain between a few hundred and a few thousand interacting cells. This range of values is, moreover, remarkably consistent with results of a recent study that used a very different method to infer the number of cells in tumour-originating niches. Across a range of tissue types, the latter study concluded that cells typically interact in communities of 300-1,900 cells.^40^

---

**Supplementary Table 1.** Characteristics of our four main models.

| Model | Gland size | Mode of cell dispersal |
| --- | --- | --- |
| Non-spatial | Effectively infinite | Not applicable |
| Gland fission | 8,192 | Glands bifurcate, such that each daughter gland inherits half of the original gland's population of cells |
| Invasive glandular | 512 | Individual cells invade neighbouring tissue and compete with normal cells |
| Boundary growth | 1 | New cells are added to the edge of the tumour |

**Supplementary Table 2.** Parameter values used in this study. Mutation rate is measured per cell division; division and dispersal rates are relative to the rates of the initial tumour cell. The effect of a driver mutation with effect size *s* is to multiply the trait value *r* by a factor of 1 + *s*(1 − *r*/*m*), where *m* is the maximum limit. Dispersal rates are set such that tumours typically take between 500 and 1,000 cell generations to grow from one to one million cells, corresponding to several years of human tumour growth.

| Parameter | Value(s) |
| --- | --- |
| Deme carrying capacity, $K$ | 1, 32, 512, 2048, 8192 |
| Driver mutation rate per cell division | $10^{-5}$ |
| Passenger mutation rate per cell division | 0.1 |
| Normal cell relative division rate | 0.9 |
| Mean value of driver effect on cell division rate | 0.1 |
| Maximum relative cell division rate | 10 |
| Maximum relative dispersal rate | 10 |
| Dispersal rate | Conditional |

**Supplementary figure 1:**
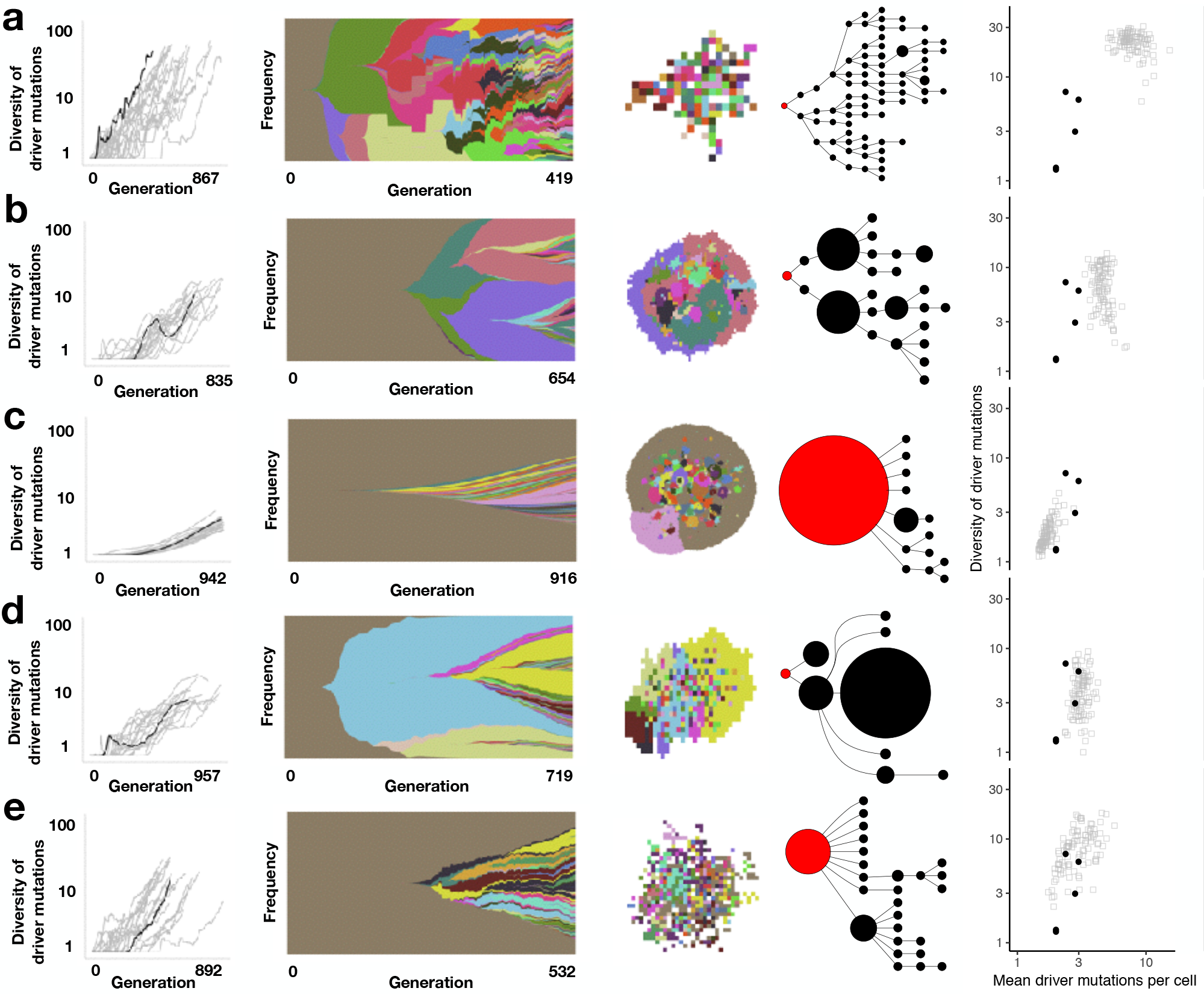
Additional evolutionary modes predicted by our model. First column: Dynamics of driver mutation diversity in 20 stochastic simulations. Diversity corresponds to the number of clones that have distinct combinations of driver mutations. A generation is defined as the expected cell cycle time of the initial tumour cell. Black curves correspond to the individual simulations illustrated in subsequent columns. These particular simulations are those with metrics closest to the medians of sets of 100 replicates. Second column: Muller plots of clonal dynamics over time. Colours represent clones with distinct combinations of driver mutations (the original clone is grey-brown; subsequent clones are coloured using a recycled palette of 26 colours). Descendant clones are shown emerging from inside their parents. Third column: Final clone proportions (for the non-spatial model) or spatial arrangement (for spatial models). For spatial models, each pixel corresponds to a patch of cells, corresponding to a tumour gland, coloured according to the most abundant clone within the patch. Fourth column: Driver phylogenetic trees. Node size corresponds to clone population size at the final time point and the founding clone is coloured red. Only clones whose descendants represent at least 1% of the final population are shown. Final column: Evolutionary metrics. Black points correspond to kidney tumour (ccRCC) data, labelled with patient codes (from ref 9). **a**, A model of tumour growth via gland fission (8,192 cells per gland), in which cells can acquire driver mutations that increase their contribution to the gland fission rate (with an average effect size of 50%), in addition to drivers that increase the cell division rate. **b**, A model in which tumour cells invade normal tissue at the tumour boundary and can also invade neighbouring glands within the tumour (512 cells per gland). **c**, A boundary-growth model of a non-glandular tumour in which cells invade neighbouring sites within the tumour. **d**, A model in which tumour cells invade normal tissue at the tumour boundary only (2,048 cells per gland). **e**, A model of tumour growth via gland fission (2,048 cells per gland). Other parameter values are listed in Supplementary Tables 1 and 2.

**Supplementary figure 2:**
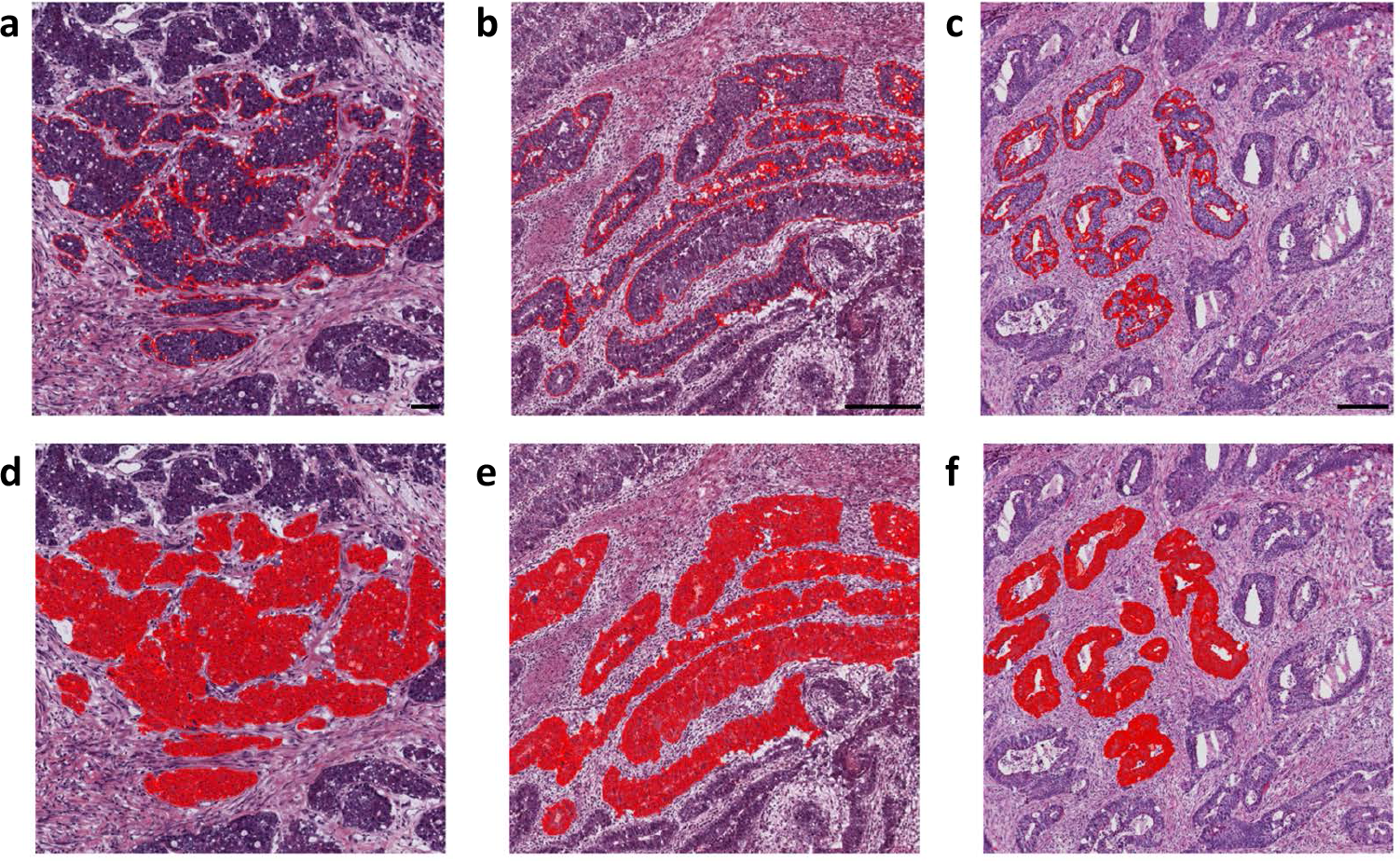
Quantification of tumour cell numbers per gland in representative patients. For three colorectal cancer patients with different sizes of tumour glands, multiple glands were manually outlined and the number of cells in each gland was counted automatically. **a**, Patient TCGA-5M-AAT6, slide 01A-01-TS1. **b**, Patient TCGA-5M-AATA, slide 01A-03-TS3. **c**, Patient TCGA-A6-2675, slide 01A-01-BS1. **d-f**, Resulting cell masks obtained with QuPath.

**Supplementary figure 3:**
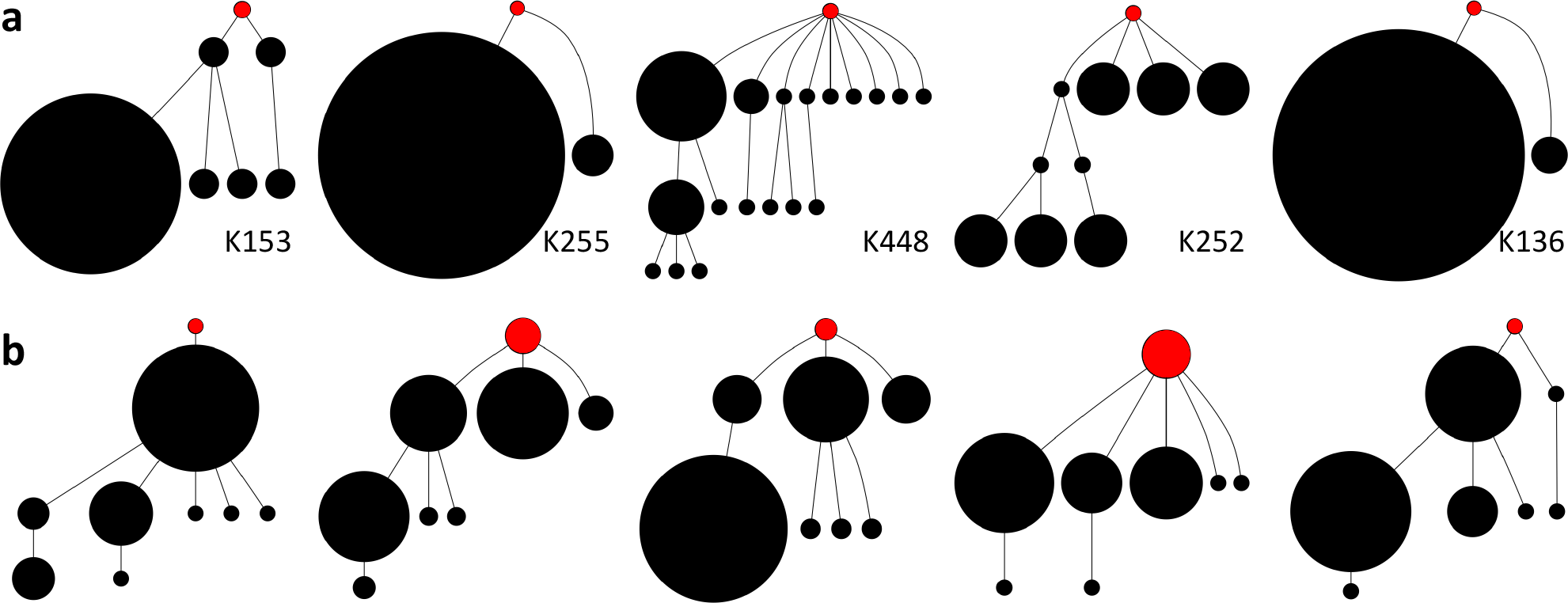
**a**, Driver phylogenetic trees for five clear cell renal cell carcinomas, labelled with patient codes. Data was obtained from data set S2 of ref 9. Clone frequencies are estimated as the proportion of regions in which the corresponding combination of driver mutations was detected. **b**, Driver phylogenetic trees resulting from an evolutionary model with tumour invasion of normal tissue at the tumour boundary (512 cells per gland; only clones whose descendants represent at least 1% of the final population are shown). Node size corresponds to clone population size. The founding clone is coloured red.

**Supplementary figure 4:**
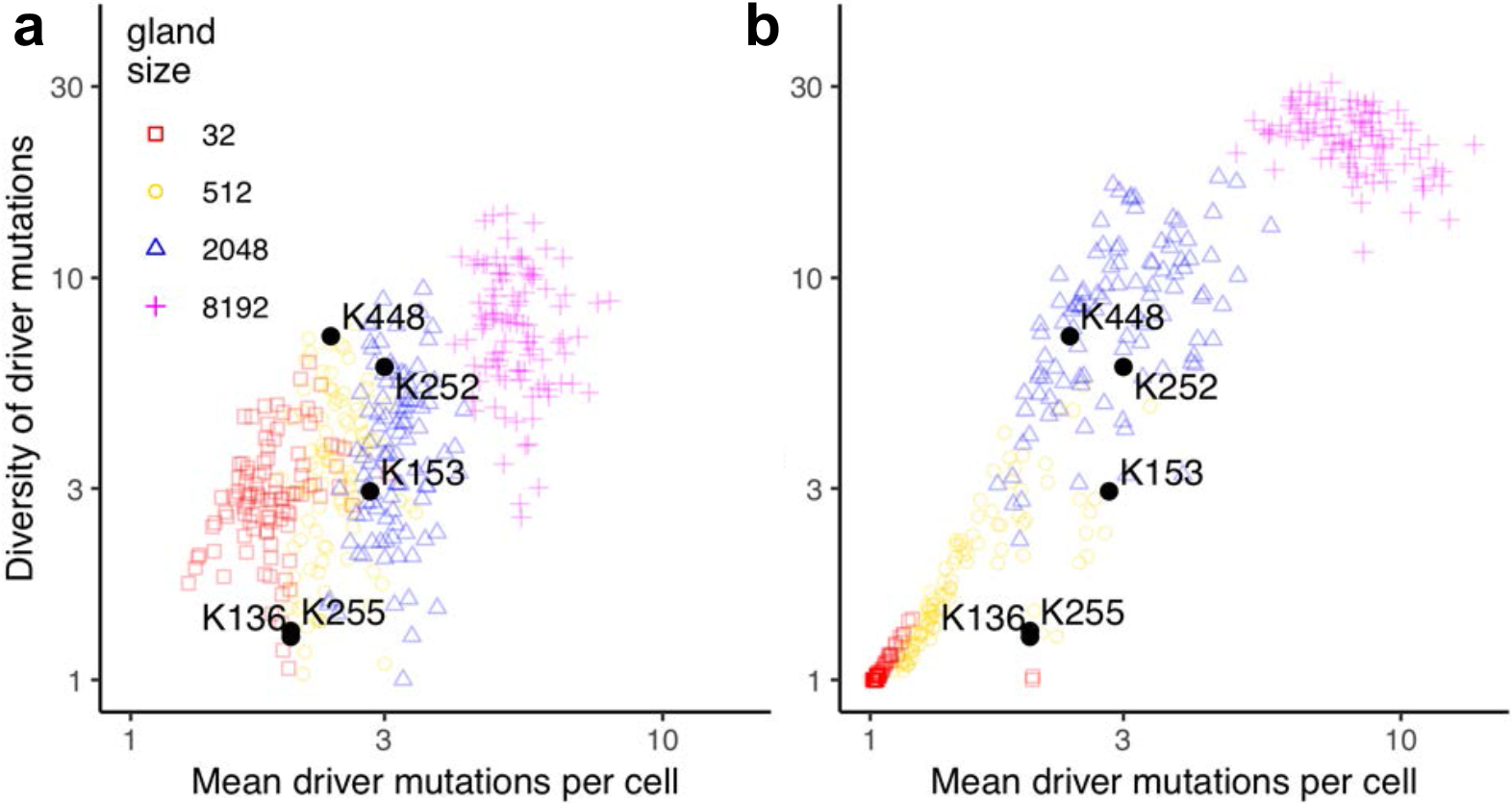
Variation in evolutionary metrics depending on gland size (100 stochastic simulations per model). **a**, Models of invasive glandular tumours. **b**, Models of tumours growing via gland fission. Black points correspond to kidney tumour (ccRCC) data, labelled with patient codes.^9^ Apart from gland size, parameter values are the same as in Figure 2.

**Supplementary figure 5:**
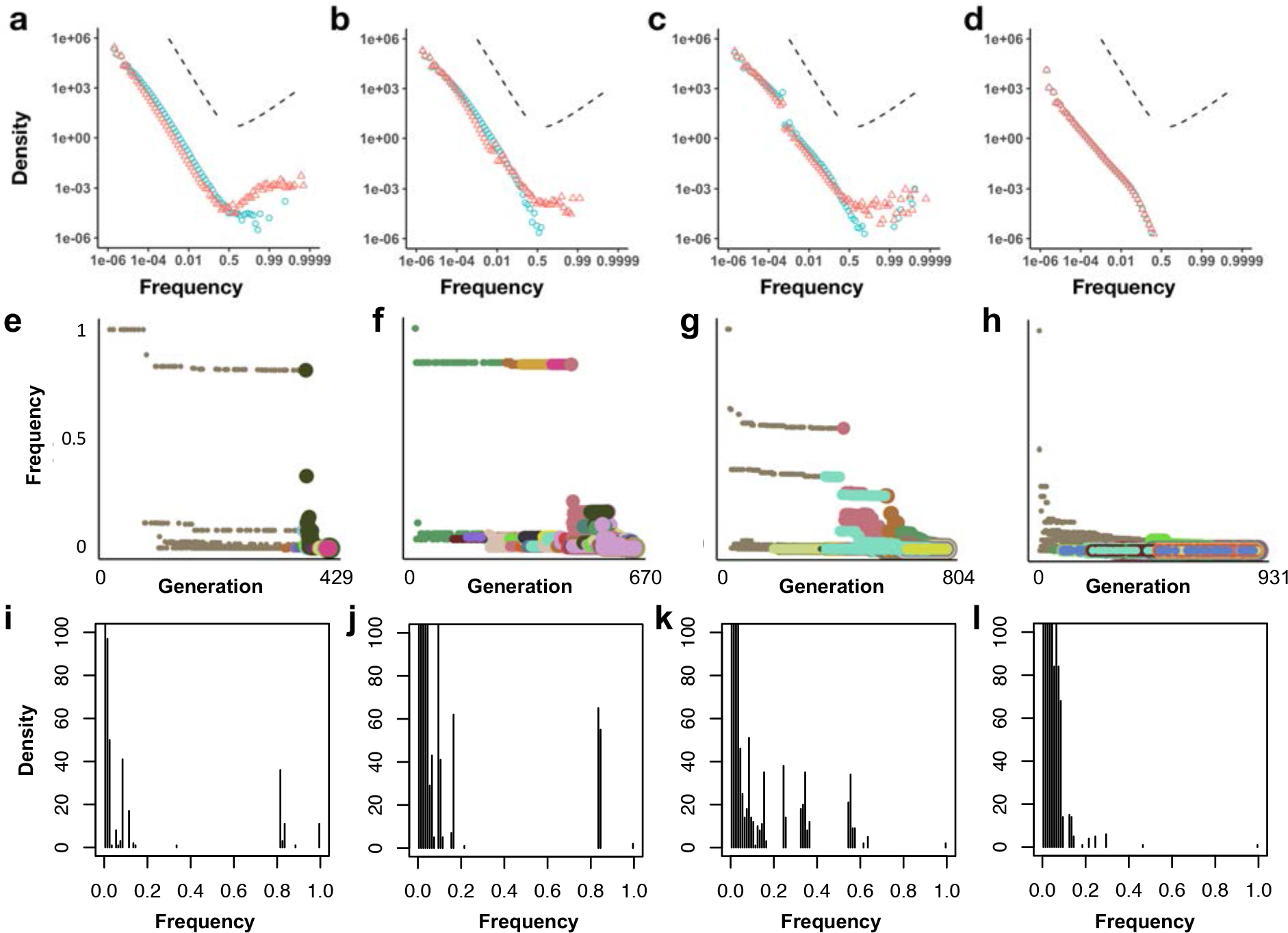
Mutation frequency distributions for simulated tumours. **a-d**, Complete mutation frequency distributions for models with only neutral mutations (blue points) or both neutral and driver mutations (red points). Each distribution represents combined data from 100 simulations of each of the four model types of Figures 2 and 3. To clarify the shape of the distributions, especially at high frequencies, the x-axes are transformed as logit(*x*) = log(*x*/(1 − *x*)), which is approximately equal to log *x* when *x* is much less than 1. Dashed lines indicate analytical predictions for an exponentially-growing population acquiring only neutral mutations (negative slope) and a population of constant size acquiring both neutral and highly beneficial mutations (positive slope^41^). **e-h**, Mutation frequency versus timing of mutation for the specific model instances of Figure 2. Point colour corresponds to clone (as in Figure 2), and size corresponds to the division rate of cells within the clone. Driver mutations are typically preceded by a string of hitchhiking passenger mutations with similar frequencies. This figure format is inspired by Figure 2 of ref 4. **i-l**, Mutation frequency distributions for the specific model instances of Figure 2, with linear axes. Results are shown for a non-spatial branching process (a, e, i); tumour growth via gland fission (b, f, j); tumour invasion of normal tissue (c, g, k); and a boundary-growth model (d, h, l). Parameter values are the same as in Figures 2 and 3.

**Supplementary Table 3.** Semi-automated quantification of tumour cell numbers per gland in histological images: raw measurements. For each region of interest (ROI) in three representative colorectal cancer patients, we measured the area of each tumour gland (in *μ*m^2^), the number of cells per gland (NumCells), the cell density in cells/mm^2^ and the distance to the nearest ROI (in *μ*m).

| Image | ROI | Area | NumCells | Cell density | Distance to nearest ROI |
| --- | --- | --- | --- | --- | --- |
| TCGA-5M-AAT6-01A-01-TS1 | ROI 1 | 4208 | 57 | 0.014 | 15 |
| TCGA-5M-AAT6-01A-01-TS1 | ROI 2 | 52263 | 588 | 0.011 | 0 |
| TCGA-5M-AAT6-01A-01-TS1 | ROI 3 | 1338 | 16 | 0.012 | 10 |
| TCGA-5M-AAT6-01A-01-TS1 | ROI 4 | 7961 | 98 | 0.012 | 0 |
| TCGA-5M-AAT6-01A-01-TS1 | ROI 5 | 19037 | 242 | 0.013 | 16 |
| TCGA-5M-AAT6-01A-01-TS1 | ROI 6 | 9832 | 131 | 0.013 | 12 |
| TCGA-5M-AAT6-01A-01-TS1 | ROI 7 | 56932 | 623 | 0.011 | 10 |
| TCGA-5M-AAT6-01A-01-TS1 | ROI 8 | 8733 | 121 | 0.014 | 0 |
| TCGA-5M-AAT6-01A-01-TS1 | ROI 9 | 33297 | 398 | 0.012 | 0 |
| TCGA-5M-AAT6-01A-01-TS1 | ROI 10 | 19531 | 241 | 0.012 | 0 |
| TCGA-5M-AAT6-01A-01-TS1 | ROI 11 | 78047 | 880 | 0.011 | 0 |
| TCGA-5M-AAT6-01A-01-TS1 | ROI 12 | 53567 | 635 | 0.012 | 0 |
| TCGA-5M-AAT6-01A-01-TS1 | ROI 13 | 3500 | 46 | 0.013 | 0 |
| TCGA-5M-AAT6-01A-01-TS1 | ROI 14 | 6479 | 83 | 0.013 | 0 |
| TCGA-5M-AAT6-01A-01-TS1 | ROI 15 | 3247 | 39 | 0.012 | 0 |
| TCGA-5M-AAT6-01A-01-TS1 | ROI 16 | 199 | 14 | 0.070 | 0 |
| TCGA-5M-AATA-01A-03-TS3 | ROI 1 | 99027 | 931 | 0.009 | 16 |
| TCGA-5M-AATA-01A-03-TS3 | ROI 2 | 33920 | 327 | 0.010 | 41 |
| TCGA-5M-AATA-01A-03-TS3 | ROI 3 | 106350 | 1056 | 0.010 | 15 |
| TCGA-5M-AATA-01A-03-TS3 | ROI 4 | 44068 | 481 | 0.011 | 0 |
| TCGA-5M-AATA-01A-03-TS3 | ROI 5 | 28618 | 298 | 0.010 | 18 |
| TCGA-5M-AATA-01A-03-TS3 | ROI 6 | 23754 | 234 | 0.010 | 0 |
| TCGA-5M-AATA-01A-03-TS3 | ROI 7 | 4586 | 53 | 0.012 | 3 |
| TCGA-5M-AATA-01A-03-TS3 | ROI 8 | 81337 | 816 | 0.010 | 0 |
| TCGA-5M-AATA-01A-03-TS3 | ROI 9 | 28697 | 249 | 0.009 | 0 |
| TCGA-5M-AATA-01A-03-TS3 | ROI 10 | 194330 | 1563 | 0.008 | 15 |
| TCGA-5M-AATA-01A-03-TS3 | ROI 11 | 10123 | 75 | 0.007 | 57 |
| TCGA-5M-AATA-01A-03-TS3 | ROI 12 | 88288 | 844 | 0.010 | 82 |
| TCGA-A6-2675-01A-01-BS1 | ROI 1 | 12145 | 136 | 0.011 | 12 |
| TCGA-A6-2675-01A-01-BS1 | ROI 2 | 6088 | 81 | 0.013 | 20 |
| TCGA-A6-2675-01A-01-BS1 | ROI 3 | 39183 | 404 | 0.010 | 19 |
| TCGA-A6-2675-01A-01-BS1 | ROI 4 | 27550 | 255 | 0.009 | 19 |
| TCGA-A6-2675-01A-01-BS1 | ROI 5 | 22363 | 217 | 0.010 | 13 |
| TCGA-A6-2675-01A-01-BS1 | ROI 6 | 23806 | 202 | 0.008 | 31 |
| TCGA-A6-2675-01A-01-BS1 | ROI 7 | 25506 | 238 | 0.009 | 0 |
| TCGA-A6-2675-01A-01-BS1 | ROI 8 | 26024 | 267 | 0.010 | 0 |
| TCGA-A6-2675-01A-01-BS1 | ROI 9 | 6718 | 74 | 0.011 | 0 |
| TCGA-A6-2675-01A-01-BS1 | ROI 10 | 9945 | 121 | 0.012 | 0 |
| TCGA-A6-2675-01A-01-BS1 | ROI 11 | 22183 | 220 | 0.010 | 0 |
| TCGA-A6-2675-01A-01-BS1 | ROI 12 | 38561 | 318 | 0.008 | 0 |
| TCGA-A6-2675-01A-01-BS1 | ROI 13 | 39801 | 306 | 0.008 | 0 |

**Supplementary Table 4.** Comparison of selected models of tumour population genetics.

| Study | Model type | Cells<br>per deme | Within-deme<br>selection? | Between-deme<br>selection? | Maximum<br>acquired<br>drivers | Exterior |
| --- | --- | --- | --- | --- | --- | --- |
| Bozic<br>*et al.* 2010<sup>25</sup> | Branching<br>process | Not<br>applicable | Not<br>applicable | Not<br>applicable | Unlimited | Void |
| Waclaw<br>*et al.* 2015<sup>19</sup> | Eden model,<br>voter model,<br>or similar | 1 | No | Yes | Unlimited | Void |
| Sottoriva<br>*et al.* 2015<sup>5</sup> | Deme fission | 10,000 | No | Yes | 1 | Void |
| Sun & Hu<br>*et al.* 2017<sup>18</sup> | Deme fission<br>(edge only) | 1,000<br>or 10,000 | Yes | No | 1 | Void |
| Current study | Any of<br>the above | Any<br>number | Yes | Yes | Unlimited | Tissue<br>or void |

